# Flux analysis reveals specific regulatory modalities of gene expression

**DOI:** 10.1101/2022.11.25.517929

**Authors:** Benjamin Martin, David Michael Suter

**Affiliations:** Institute of Bioengineering, School of Life Sciences, Ecole Polytechnique Fédérale de Lausanne (EPFL), Lausanne 1015, Switzerland

**Author notes:** **For correspondence:** (BM); (DMS).

## Abstract

The level of a given protein is determined by rates of mRNA and protein synthesis and degradation. The reliable quantification of these rates is thus essential to understand how protein levels are regulated. While several studies have quantified the contribution of different steps of gene expression in regulating protein levels, these are limited by the use of equilibrium approximations when studying out-of-equilibrium biological systems. Here we introduce flux analysis to quantify the contribution of gene expression steps to regulate the dynamics of the expression level for specific proteins. We use flux analysis to analyze a published RNA-seq and proteomics dataset of mouse bone marrow-derived DCs stimulated with LPS. Our analysis revealed six regulatory modalities shared by sets of genes with clear functional signatures. We also find that protein degradation plays a stronger role than expected in the adaptation of protein levels to LPS stimulation. These findings suggest that shared regulatory strategies can lead to versatile responses at the protein level and highlight the importance of going beyond steady-state approximations to uncover the dynamic contribution of different steps of gene expression to protein levels.

## Introduction

Proteins are the main workhorses of cells and thus ultimately define their function. As end-products of the gene expression cascade, protein levels are ultimately governed by rates of transcription, RNA decay, translation, and protein decay. While in principle there is a large combinatorial space for gene expression rate changes to reach a desired protein level, these rates are constrained by several factors. Genes with high transcription and low translation are depleted as predicted by considering a trade-off between precision (to reduce stochastic variation in protein number) and economy (to reduce the energetic cost of mRNA synthesis) (***Hausser et al., 2019***). Gene expression rates also display a strong tissue-specificity. For instance, RNA degradation rate is on average lower in the heart than in the liver (***Hammond et al., 2016***). Gene expression rates can also be constrained by the need for rapid changes in protein levels (***Signer et al., 2014***; ***Wang et al., 2018***). For example, in the liver the high RNA degradation rate of the gluconeogenic gene G6pc has been shown to allow rapid reduction of both its RNA and protein levels upon refeeding, which is central to shut down hepatic glucose output as fast as possible (***Bahar Halpern et al., 2015***).

Which step(s) of gene expression dominate in regulating differences in the expression level of different proteins has been debated over the past decade (***Buccitelli and Selbach, 2020***). Some studies argued for the central role of translation and protein degradation (***Brion et al., 2020***; ***Ghazalpour et al., 2011***; ***Kristensen et al., 2013***; ***Schwanhäusser et al., 2011***; ***Zhang et al., 2014***), while others concluded that RNA level makes a larger contribution to determine protein levels (***Battle et al., 2015***; ***Jovanovic et al., 2015***; ***Li et al., 2014***; ***Li and Biggin, 2015***).

Another unsolved question is how different steps in gene expression are modulated to change the expression level of a given protein, for instance upon cell stimulation. In the context of dynamic changes in proteome content through external stimulation, ***Jovanovic et al. (2015***) quantified which gene expression parameter(s) explain most protein level variance on a genome-wide scale. They concluded that RNA levels explain 43% of absolute changes in protein expression (***Jovanovic et al., 2015***). However, this study is limited by the fact that observed vs. expected protein levels are computed under the steady-state assumption in a system that is out-of-equilibrium (***Martin and Suter, 2022***). The reliable quantification of dynamic changes in rates of different gene expression steps (***Alber et al., 2018***; ***Elowitz et al., 2002***; ***Hilfinger and Paulsson, 2011***) in out-of-equilibrium systems remains a central challenge.

In this study, we introduce flux analysis to quantify the contributions of different regulatory steps of gene expression in determining protein levels and proteome dynamics upon perturbation, both proteome-wide and for specific proteins. We uncover an unexpected weight for protein degradation in regulating gene expression dynamics and uncover a limited number of regulatory modalities that are shared by sets of genes with clear functional signatures.

## Results

### Protein degradation rate is a major determinant of protein levels and proteome dynamics upon perturbation

Here we focus on a study that attempted to determine the weight of the different steps of gene expression on changes in protein levels in dendritic cells (DCs) stimulated with Lipopolysaccharide (LPS) (***Jovanovic et al., 2015***). RNA sequencing and pulsed-SILAC were performed to quantify changes in RNA and protein levels at different time points, which was used to build a quantitative model of gene expression and to infer rate values of the model for all proteins detected by SILAC. The authors concluded that changes in transcriptome and proteome upon LPS stimulation of DCs are mainly driven by changes in RNA levels (***Jovanovic et al., 2015***).

This study used a steady-state approximation, which underlies their linear correlation analysis based on Pearson’s correlation coefficient. We reasoned that this assumption may limit the reliability of gene expression rate inferences when gene expression is abruptly altered by external stimulation. We thus decided to develop a new approach allowing us to infer gene expression states away from equilibrium. Our strategy is based on the decomposition of the protein flux in its different components (Methods and Materials) and allows us to quantify the relative contributions of each regulatory process, i.e., RNA level, translation, protein level, and degradation (***Figure 1***A) to changes in proteome composition upon external stimulation.

**Figure 1.**
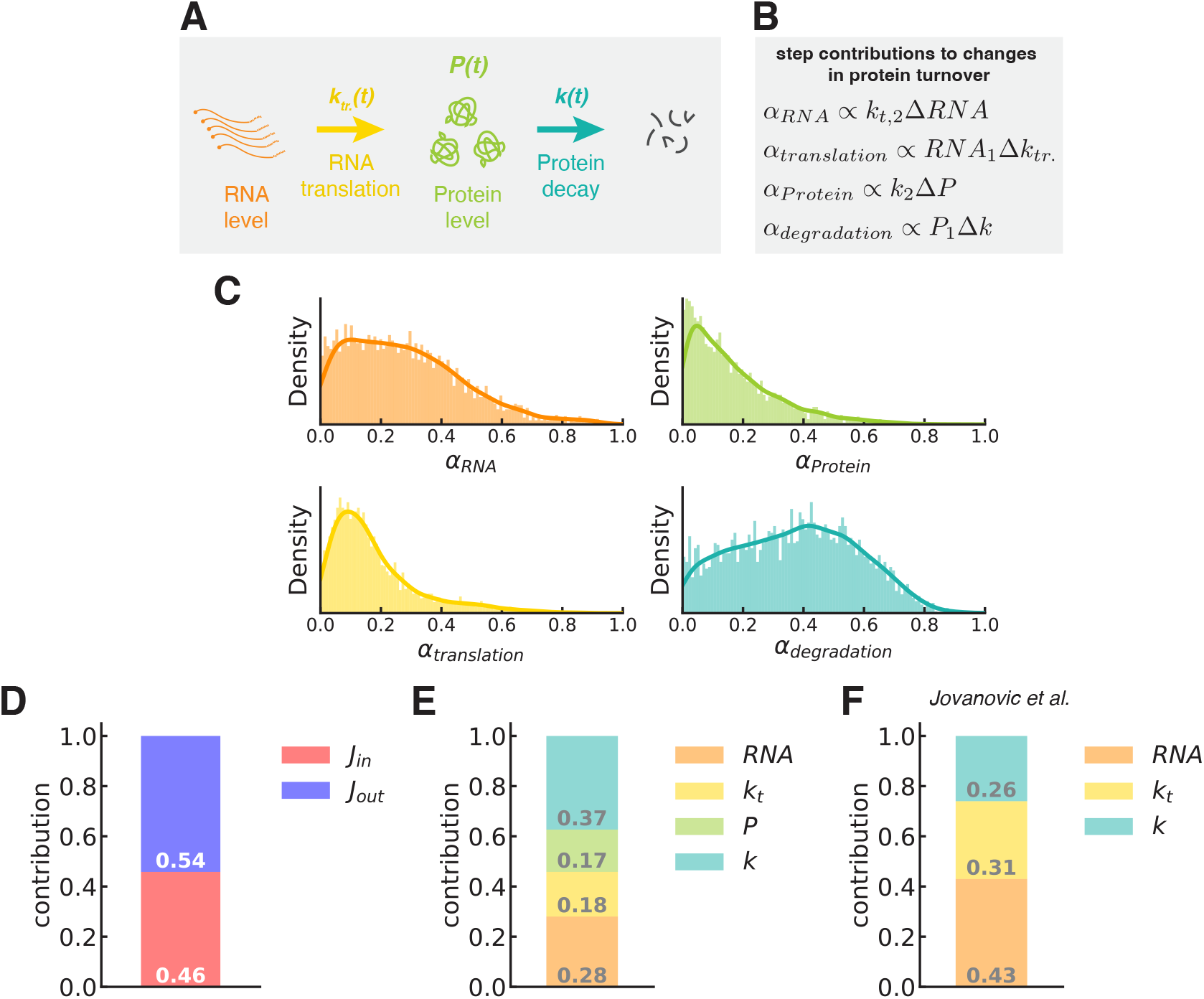
Protein degradation is the main contributor to proteome recomposition upon DCs stimulation with LPS. **A**. A gene expression system can be decomposed into four distinct regulatory steps — RNA level, RNA translation, Protein level, and Protein decay, each of which alter the protein flux in the system. **B**. The contribution of each step can be quantified using the rate values — translation rate *k*_*tr*_ and degradation rate *k* — along with the RNA and protein (*P*) levels. **C**. Distribution of the contributions of the different regulatory steps for the set of proteins studied in Jovanovic et al. (Jovanovic et al., 2015). The plain lines represent the kernel density estimations from the associated histograms, for visualization purposes. **D**. Barplot of the mean contribution of the normalized influx ‖ Δ*J*_*in*_ ‖ (*J*_*in*_) and outflux ‖ Δ*J*_*out*_ ‖ (*J*_*out*_) for the entire set of proteins studied. **E**. Barplot of the mean contribution of the different regulatory steps for the entire set of proteins studied. **F**. Contributions to the absolute change in protein abundances after LPS stimulation found in ***Jovanovic et al. (2015***). **Figure 1—figure supplement 1**. Contributions to protein flux can be computed in a time-dependent manner even out-of-equilibrium. **Figure 1—figure supplement 2**. RNA level and protein degradation are the main contributor to protein flux changes for protein-level upregulation and downregulation, respectively. **Figure 1—figure supplement 3**. Change in protein flux for upregulated immune response proteins is mainly driven by RNA level changes. **Figure 1—figure supplement 4**. Immune response proteins exhibit variable contributions of the different regulatory steps. **Figure 1—figure supplement 5**. Change in protein flux for mitochondrial proteins is mainly driven by protein degradation. **Figure 1—figure supplement 6**. Increase in degradative flow of mitochondrial proteins leads to the metabolic rewiring of DCs after LPS stimulation. **Figure 1—figure supplement 7**. Translation is a key regulatory steps in protein flux changes of ribosomal proteins. **Figure 1—source data 1**. https://github.com/UPSUTER/FluxAnalysis/Figure1.ipynb **Figure 1—source code 1**. https://github.com/UPSUTER/FluxAnalysis/data from ***Jovanovic et al. (2015***)

Briefly, we defined the total flux in the gene expression system in a state *i* as the sum of the influx and outflux, namely ‖ *J*_*tot,i*_ ‖=‖ *J*_*in,i*_ ‖ + ‖ *J*_*out,i*_ ‖ (Methods and Materials). *J*_*in,i*_ = *s*_*i*_ and *J*_*out,i*_ = *k*_*i*_ × *P*_*i*_,*s*_*i*_ and *k*_*i*_ being the protein synthesis and degradation rates, respectively, and *P*_*i*_ the protein level in state *i* (*i* = 1 for initial state and *i* = 2 for the final state in ***Figure 1***B). We derived the influx and outflux contributions, *α*_*in*_ and *α*_*out*_, as the normalized changes in influx ‖ Δ*J*_*in*_ ‖ / ‖ Δ*J*_*tot*_ ‖ and outflux ‖ Δ*J*_*out*_ ‖ / ‖ Δ*J*_*tot*_ ‖ with respect to the total flow ‖ Δ*J*_*tot*_ ‖. We then decomposed *α*_*in*_ in *α*_*RNA*_ and *α*_*translation*_ ; and *α*_*out*_ in *α*_*prot*_ and *α*_*deg*_. They represent the contribution of the RNA level, the translation rate, the protein level, and the degradation rate to the total variation in flux (Methods and Materials, ***Figure 1***A and 1B), respectively. Note that the contribution *α*_*prot*_ of the protein level to the outflux contribution *α*_*out*_ comes from the fact that we considered the protein degradation process as a first-order reaction. The contribution of the variation in influx *α*_*in*_ and outflux *α*_*out*_ (Methods and Materials) as well as the contribution of the different regulatory steps to the total variation in flux, can be computed in a time-dependent manner and away from equilibrium (***Figure 1—figure Supplement 1***A and ***Figure 1—figure Supplement 1***B).

We first used flux analysis to ask how different steps of gene expression contribute to changes in protein levels upon LPS stimulation. We found that different protein expression systems show variable contributions of the different components, suggesting different regulatory modalities of gene expression upon perturbation (***Figure 1***C). The most variable contributions are the ones linked to RNA level and protein degradation, highlighting the central role of these two steps in mediating changes in protein turnover and proteome composition induced by LPS stimulation (***Figure 1***C). This partly recapitulates ***Jovanovic et al. (2015***) results obtained through variance analysis, i.e., Pearson’s correlation coefficient decompositions (***Jovanovic et al., 2015***). Influx ‖ Δ*J*_*in*_ ‖ and outflux ‖ Δ*J*_*out*_ ‖ alterations nearly equally contribute to the total alteration of protein flux in gene expression systems after stimulation of DCs with LPS (***Figure 1***D). Protein degradation is the main contributor (37% on average) of proteome re-composition dynamics upon DCs stimulation (***Figure 1***E). It is followed by RNA level (28%). Translation and protein level provide a smaller contribution to changes in the dynamics of protein expression (***Figure 1***E). In summary, we find that unexpectedly, modulation of protein degradation rates plays a broader role than mRNA abundance dynamics in determining protein levels dynamics upon LPS stimulation proteome-wide.

We next asked how different steps of gene expression contributed specifically to the up- or downregulation of protein levels. In line with ***Jovanovic et al. (2015***), mRNA levels and translation efficiency contributed more substantially to the upregulation of protein levels (***Figure 1—figure Supplement 2***A), whereas changes in protein degradation mostly contributed to the downregulation of protein levels (***Figure 1—figure Supplement 2***B). Similarly, we found that flux modulation in upregulated immune response proteins (***Rosenfeld et al., 2018***) was dominated by RNA level contributions (***Figure 1—figure Supplement 3***). For example, as in ***Jovanovic et al. (2015***), RNA level is the main contributor to changes in protein levels for the negative immune regulator Trafd1 (***Figure 1—figure Supplement 4***). We then focused on Cebpb and Rela, two key regulators of DCs and of the LPS response (***Jovanovic et al., 2015***). While Rela changes in response to LPS were mainly driven by changes in protein degradation rate, those of Cebpb were mainly driven by changes in protein level *P* (***Figure 1—figure Supplement 4***). While ***Jovanovic et al. (2015***) reported that Rela protein level is primarily regulated at the level of translation and protein degradation, here again, we find a stronger contribution of the degradation process in Rela gene expression response to LPS stimulation.

We were next interested to focus on mitochondrial proteins since DCs show mitochondrial metabolic rewiring upon LPS stimulation. Using MitoCart3.0 (***Rath et al., 2021***), we found 364 mitochondrial proteins in the ***Jovanovic et al. (2015***) dataset (***Figure 1—figure Supplement 5***A). We found that protein degradation is the main contributor to mitochondrial protein expression alteration after LPS stimulation (Supplementary Figures ***Figure 1—figure Supplement 5***B and ***Figure 1—figure Supplement 5***C), in line with ***Jovanovic et al. (2015***). The degradation-mediated flux increased on average more (1.2-fold) for mitochondrial proteins than for all other proteins (***Figure 1—figure Supplement 6***A). As already suggested by the approach of ***Jovanovic et al. (2015***), protein degradation explains almost all protein flux changes of key mitochondrial enzymes such as Sucla2, Aco2, and Suclg1 (***Figure 1—figure Supplement 6***B). This observation is consistent with a shift from oxidative phosphorylation and oxygen consumption to glycolysis, glucose consumption, and lactate production in DCs stimulated with LPS (***Everts et al., 2014, 2012***; ***Jovanovic et al., 2015***; ***Krawczyk et al., 2010***; ***Pearce and Pearce, 2013***). Similarly, we found that ribosomal protein (***Tweedie et al., 2021***) expression is mainly regulated through translation and protein degradation alteration (58% in total, ***Figure 1—figure Supplement 7***A-C) in response to LPS treatment. Finally, translation-mediated flux increased on average more (1.75-fold) for ribosomal proteins than for all other proteins (***Figure 1—figure Supplement 7***D).

Taken together, our results suggest that protein degradation is a key genome-wide step of protein flux modulation in response to DC stimulation with LPS.

### Flux analysis distinguishes between different regulatory modes of gene expression

We next aimed to use flux analysis to unravel different modalities of dynamic regulation of gene expression upon cell perturbation. To do so, we clustered genes according to their contributions *α*s to the gene-specific total flux into six clusters (Figures 2A-B and ***Figure 2—figure Supplement 1***). Clusters A and E exhibit a higher contribution of the influx, ‖ Δ*J*_*in*_ ‖, clusters C, D, and F a higher contribution of the outflux, ‖ Δ*J*_*out*_ ‖, and cluster B a similar contribution of both fluxes. The total flux, i.e., protein flux, is mainly affected by variation of RNA levels in cluster A. In cluster B, the main contribution comes from both RNA level and protein degradation. In cluster C, the main contribution comes from both protein level and protein degradation. In clusters D, E, and F, protein degradation, translation, and protein level, respectively, are the main contributors to changes in protein flux. We then split each cluster into two subclusters according to up- vs down-regulation of the fluxes for the different regulatory steps of protein expression (***Figure 2***C-D). This analysis suggests that the different regulatory modes we found can each lead to diverse changes in protein flux, RNA levels, and protein levels (***Figure 2***C-D and ***Figure 2—figure Supplement 2***). In other words, a given regulatory modality can underly opposite protein level dynamics.

**Figure 2.**
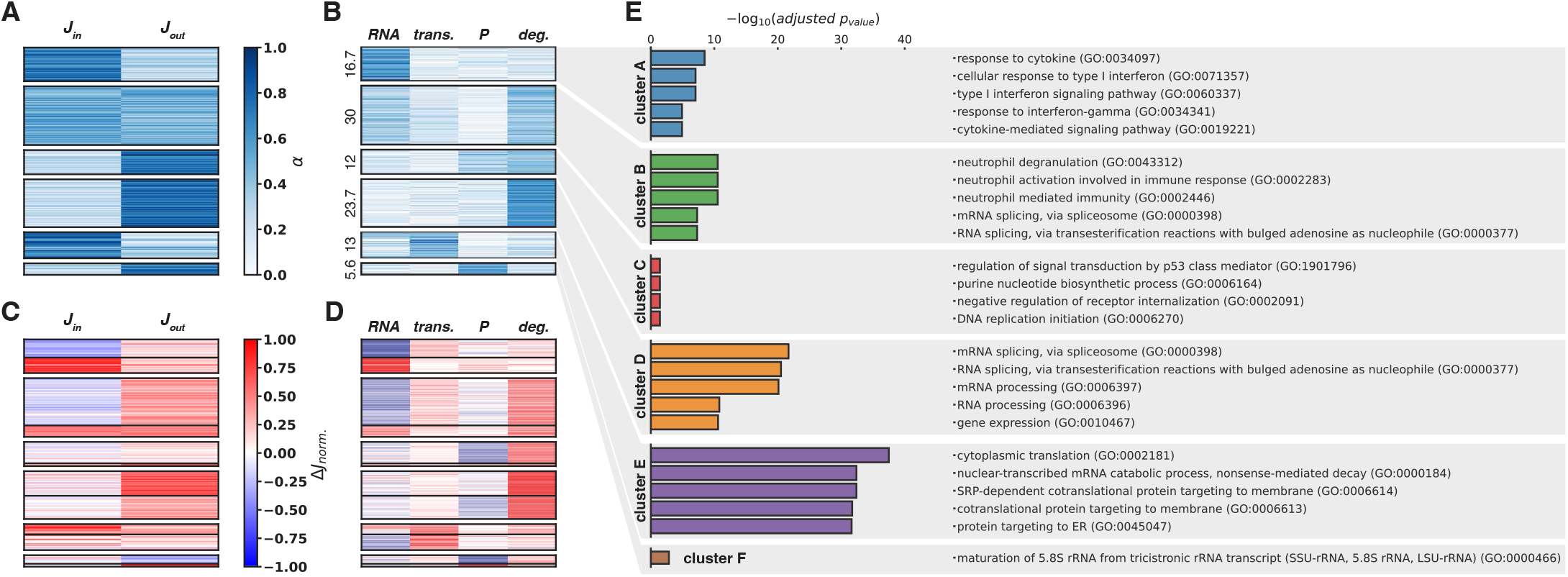
Six regulatory modalities of protein flows can be identified. **A**. and **B**. Hierarchical clustering on step contributions identifies six main regulatory modalities (clusters A to F) of protein flows. **C**. and **D**. Each cluster can be decomposed into two subclusters according to the algebraic values of the step contributions, i.e., the normalized flux change denoted as Δ*J* _*norm*_ (Methods and Materials). **E**. GO (Biological Processes 2021) terms enrichment analysis on the six clusters identified in **A-D**. *trans*. stands for translation, *P* for protein levels. **Figure 2—figure supplement 1**. Gap statistics suggest the presence of six clusters in our dataset. **Figure 2—figure supplement 2**. Regulatory modes are linked to versatile changes in protein flux, RNA, and protein levels. **Figure 2—figure supplement 3**. GO Biological processes terms can be clustered in six clusters of regulatory modalities.. **Figure 2—source data 1**. https://github.com/UPSUTER/FluxAnalysis/Figure2.ipynb **Figure 2—source code 1**. https://github.com/UPSUTER/FluxAnalysis/data from ***Jovanovic et al. (2015***)

Finally, we performed gene ontology terms enrichment analysis (***Chen et al., 2013***; ***Fang et al., 2022***; ***Mootha et al., 2003***; ***Subramanian et al., 2005***) to decipher if the regulatory modalities we found were preferentially related to specific biological functions (***Figure 2***E and ***Figure 2—figure Supplement 3***). Our findings were largely consistent with those previously reported (***Jovanovic et al., 2015***). Briefly, acute response genes to LPS stimulation are over-represented in cluster A, and gene expression-related genes are mostly over-represented in clusters D and E (***Figure 2***E and ***Figure 2—figure Supplement 3***). Cluster A genes that are enriched in stimulus-specific genes are mainly (an average of 57%) regulated through RNA level, for both up and downregulation (***Figure 2***D). Conversely, genes related to transcription, RNA metabolism, and translation are primarily regulated by modulation of translation and protein degradation (Figures 2D and 2E. We also found that highly expressed proteins (***Figure 2—figure Supplement 2***), identifiable to cluster E and including ribosomal proteins (***Figure 2***E and ***Figure 2—figure Supplement 3***), are mainly regulated through translation. In summary, we show that flux analysis allows us to determine six primary modalities for gene expression regulation upon cell perturbation with consistent functional signatures.

## Discussion

In this brief report, we present a new mathematical framework to quantify the contribution of regulatory steps to gene expression regulation upon perturbation. Using this new quantitative method and a classical cellular stimulation model (***Jovanovic et al., 2015***), we questioned the notion that states that most dynamic changes in protein levels emanate from changes in RNA levels. We find that protein degradation is a strong, proteome-wide determinant of protein level variations (***Raj et al., 2006***) and proteome dynamics upon perturbation. We identified six primary modalities for gene expression regulation upon cell perturbation, which represents only a subset of 16 possible regulatory modalities when assuming steps as independent binary switches. The six clusters we found exhibit clear functional signatures in our system. This limited number of regulatory modalities can either indicate that LPS does not unravel all the possible regulation modalities of gene expression or that these modalities are tightly constrained

While upregulation of IFN-regulated genes (mainly found in cluster A) can be explained by an increase in RNA levels due to the presence of interferon-sensitive response element (ISRE) in their upstream regulatory sequences, the molecular mechanisms underlying the five other regulatory strategies remain to be elucidated. Specific DNA, RNA, protein sequence, or structural motifs may be specifically present in the different clusters. For instance, it has been shown that some RNA coding for protein biosynthesis factors exhibit a 5’ Terminal OligoPyrimidine (5’ TOP) motif (***Cockman et al., 2020***). Under stress conditions such as amino-acids starvation or hypoxia, 5’TOP mRNAs are subjected to translation repression, polysome release, and accumulation in stress granules (***Cockman et al., 2020***; ***Damgaard and Lykke-Andersen, 2011***).

The generalizability of our conclusions is mainly limited by the availability of datasets akin to ***Jovanovic et al. (2015***). Therefore, whether the different regulatory modes we identified can be found in other contexts remains an open question. Another limitation is the current lack of information on transcriptome dynamics. In the near future, this could be overcome by combining the newly developed mass-spectrometry (***Brunner et al., 2022***; ***Fulcher et al., 2022***; ***Leduc et al., 2022***; ***Slavov, 2022***), metabolic labeling and sequencing methods (***Erhard et al., 2019***; ***Herzog et al., 2017***). RNA dynamics is especially important in our context. It has been suggested that RNA degradation is important for controlling drastic changes in gene expression in immune cells subject to stimulations (***Ivanov and Anderson, 2013***). For instance, in DCs stimulated with LPS, it has been proved that mRNAs of house-keeping genes are degraded slowly while those of genes involved in transcriptional regulation, immune signaling, and those of genes encoding cytokines are degraded quickly (***Kumagai et al., 2016***; ***Yang et al., 2003***). More broadly, post-transcriptional regulation of mRNA metabolism has been proven a key determinant of gene expression in different biological processes (***Franks et al., 2017***; ***Hansen et al., 2018a***,b; ***Matkovic et al., 2022***) and should therefore not be eluded. Our analytical framework should be broadly applicable to study changes in gene expression fluxes and gene expression modalities in contexts where simplifying assumptions such as equilibrium and/or linear time variations in rates break down.

## Methods and Materials

### Model of protein expression

We assume that the dynamics of protein level *P* can be described by a simple ordinary differential equation (***Figure 1***):

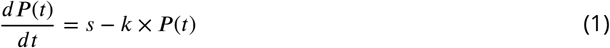

in which *s* and *k* are, respectively, the production and decay rates of the protein. This equation is readily solvable for *s* and *k* constants. The dynamic of *P* follows a canonical exponential relaxation dynamic (assuming *P* (*t* = 0) = 0):

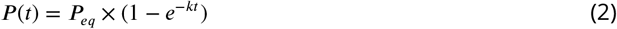

where the level of protein at equilibrium *P*_*eq*_ is given by:

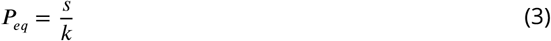

To illustrate the time dependency of *s* and *k* in the governing ODE we can rewrite ***Equation 1***:

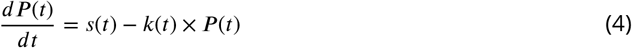

We can then define the influx *J*_*in,t*_ and outflux *J*_*out,t*_ at time *t* as:

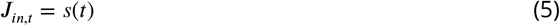

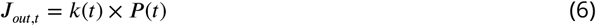

Importantly, the protein synthesis rate *s* depends on the RNA level *RNA* and the translation rate *k*_*tr*._ (***Figure 1***).

### Decoupling fluxes in regulation steps components

#### Principle

We can decompose the flux in the gene expression system in its different regulatory — biochemical — steps. To do so, let us define:

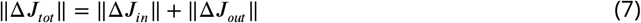

in which Δ*J*_*tot*_ is the total variation in flux, between two states (1 and 2), and Δ*J*_*in*/*out*_ is the variation in influx and outflux respectively. ||.|| denotes here the sum of the absolute value of the flux components (see after). We define the (normalized) contributions *α*_*in*/*out*_ to the total variation in flux as:

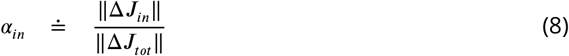

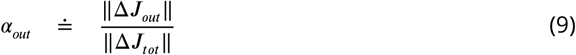

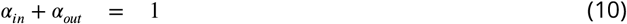

We can even decompose these two quantities as:

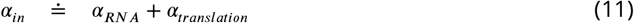

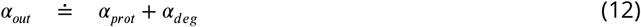

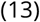

with:

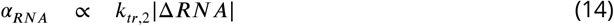

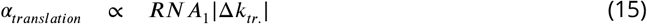

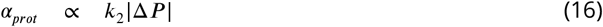

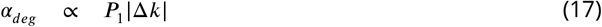

*α*_*RNA*_, *α*_*translation*_, *α*_*prot*_, *α*_*deg*_ represent, respectively, the contribution of the RNA level, the translation rate, the protein level, and the degradation rate to the total variation in flux as defined previously. 1 and 2 subscripts specify the state. For example, *P*_1_ means the protein level in state 1. Δ is the difference between the two states. It is worth noting that *α* ∈ [0, 1].

We also defined Δ*J*_*norm*_, which represents the algebraic (normalized) contribution to changes in the total flux. For instance for the RNA step, for a transition between states 1 and 2, we have:

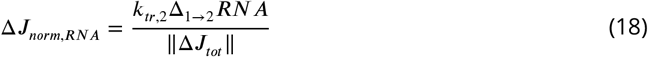

It is worth noting that Δ*J*_*norm*_ ∈ [−1, 1].

#### Derivation

Here we give detailed steps to derive the contributions *α*s. Let’s start looking at Δ*J*_*in*_:

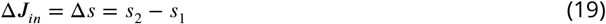

Since, by definition *s*(*t*) = *k*_*tr*._(*t*) × *RNA*(*t*):

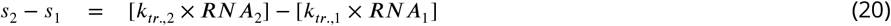

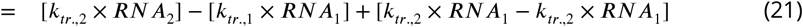

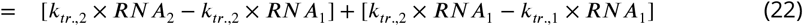

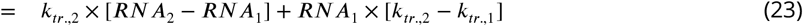

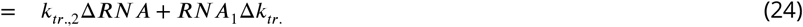

To summarize we have:

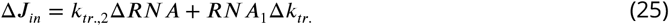

Using the exact same decomposition principle we have:

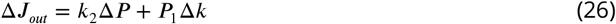

We observe this decomposition allows us to distinguish variations in flux mediated by change in one and only one component of the system. Because we are interested in the amplitude of these variations, i.e. in their singular absolute values, and not directly their algebraic values and compensatory nature, we define the operator ‖.‖ such that:

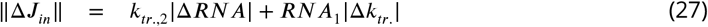

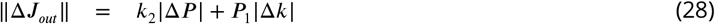

And finally, the total variation in flux intuitively reads:

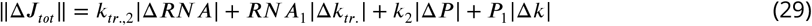

The components *α*s are extracted from these previous expressions:

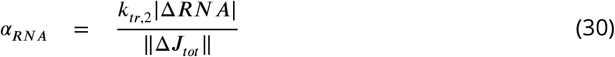

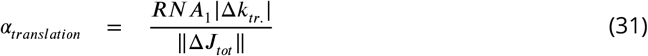

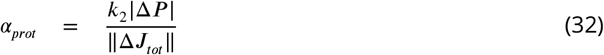

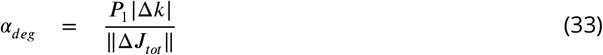

It follows that by definition:

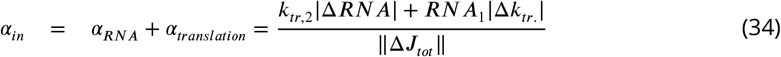

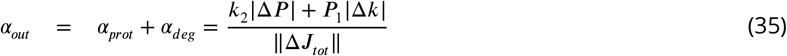

That can be rewritten:

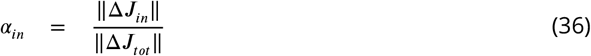

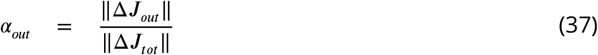

#### Computation

In this subsection, we give an example of how we compute the contributions for one gene expression system (“gene”/”protein”) using Jovanovic et al. data. To compute the contributions *α*s, we need the translation rates, the protein degradation rates, the RNA levels, and protein levels in states 1 (0h LPS stimulation) and 2 (12h LPS stimulation). For Tradf1 these values are given in Table 1.

**Table 1.**

| $i$ | 0h (state 1) | 12h (state 2) |
| --- | --- | --- |
| $k_{tr,i} (IMS.TPM^{-1}.h^{-1})$ | 0.012 | 0.021 |
| $RNA_i (TPM)$ | 80 | 298 |
| $k_i (h^{-1})$ | 0.087 | 0.094 |
| $P_i (IMS)$ | 23.9 | 54.2 |

From this, we can compute the algebraic variations:

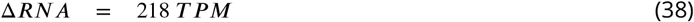

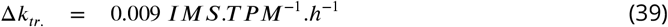

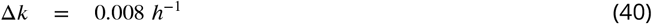

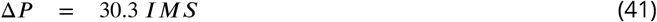

That leads to the computation of the total variation in flux:

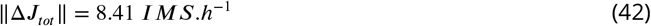

With all the previous intermediate computations we can finally compute the contributions *α*s:

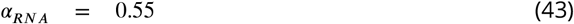

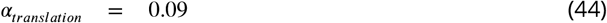

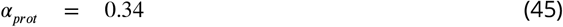

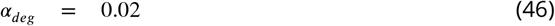

### Datasets

We employed proteomics and transcriptomics measurements from ***Jovanovic et al. (2015***). The rates — translation and protein degradation — were directly taken from the modeling and fitting done in the original paper (***Jovanovic et al., 2015***). We averaged all the measurements and rates over the two replicates of LPS conditions (0h and 12h) performed in the study.

### Gene ontology terms enrichment analysis

Gene ontology (GO) terms enrichment analysis was performed using `GSEApy` (***Fang et al., 2022***) in Python 3.9.13. Further details are given in the legends of the figures.

### Data clustering

Data were clustered using agglomerative clustering with Euclidian affinity and Ward linkage. We used the `AgglomerativeClustering` function from the scikit-klearn 1.1.1 (***Pedregosa et al., 2011***) module in Python 3.9.13. The number of clusters was determined using gap statistics (***Tibshirani et al., 2001***) computation using the `gap_statistic` package in Python 3.9.13 (***Granger, 2022***) with in-house modifications.

## Acknowledgments

We thank Maike Hansen, Félix Naef, Paolo de Los Rios, and Almut Eisele for their comments on an earlier version of this manuscript. We thank Maike Hansen, Sun Shoujie, Joanna Dembska and Félix Naef for thoughtful discussions and suggestions.

## Author contributions

B.M. conceptualized the project and analyzed the data. B.M. and D.S. interpreted the data and wrote the manuscript.

---

**Figure 1—figure supplement 1.**
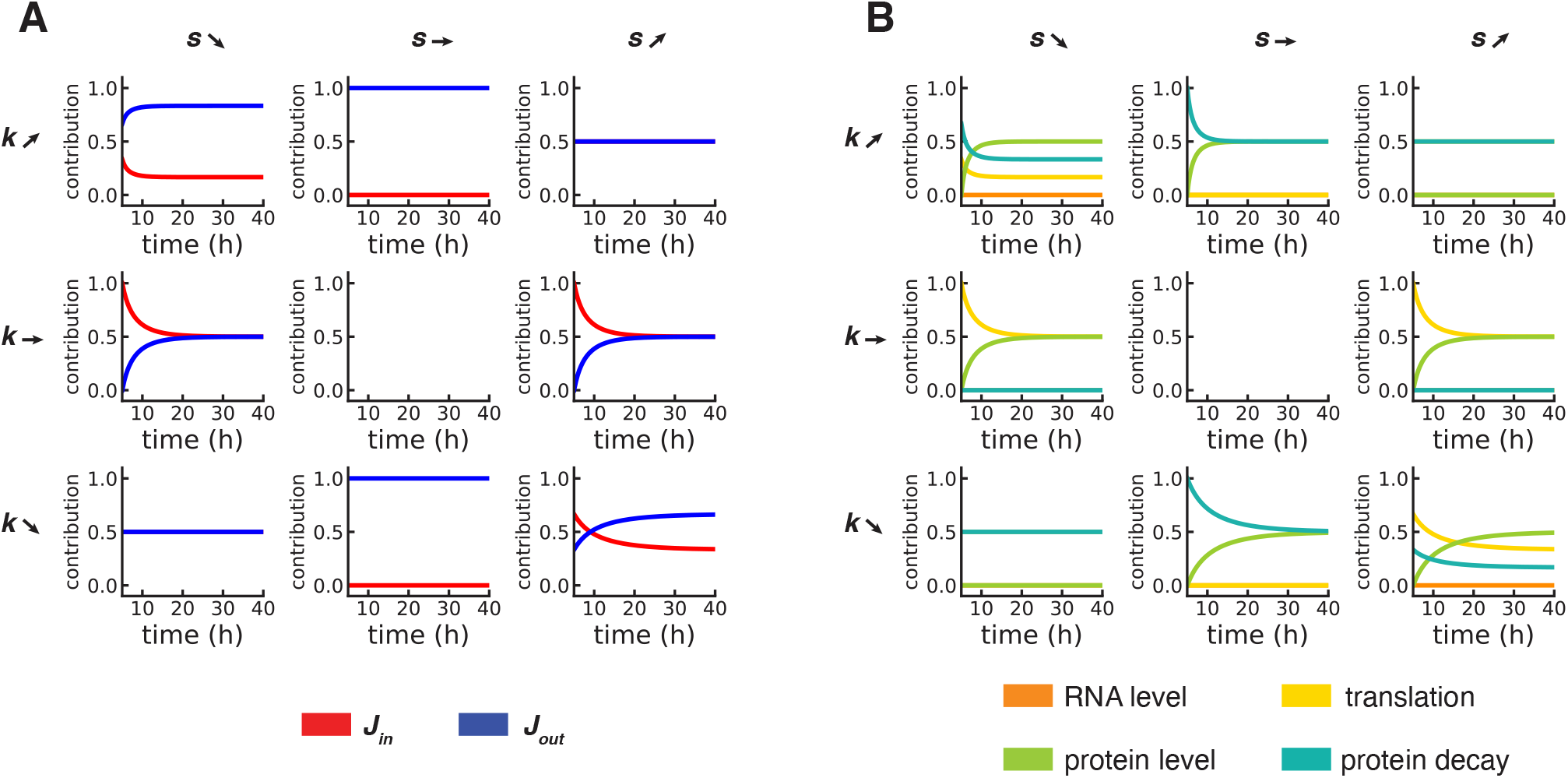
Contributions to protein flux can be computed in a time-dependent manner even out-of-equilibrium. **A**. Time-dependent contribution of the influx and outflux in the same setup as presented in ***Figure 1*. B**. Time-dependent contribution of the regulatory steps in the same setup as presented in ***Figure 1***.

**Figure 1—figure supplement 2.**
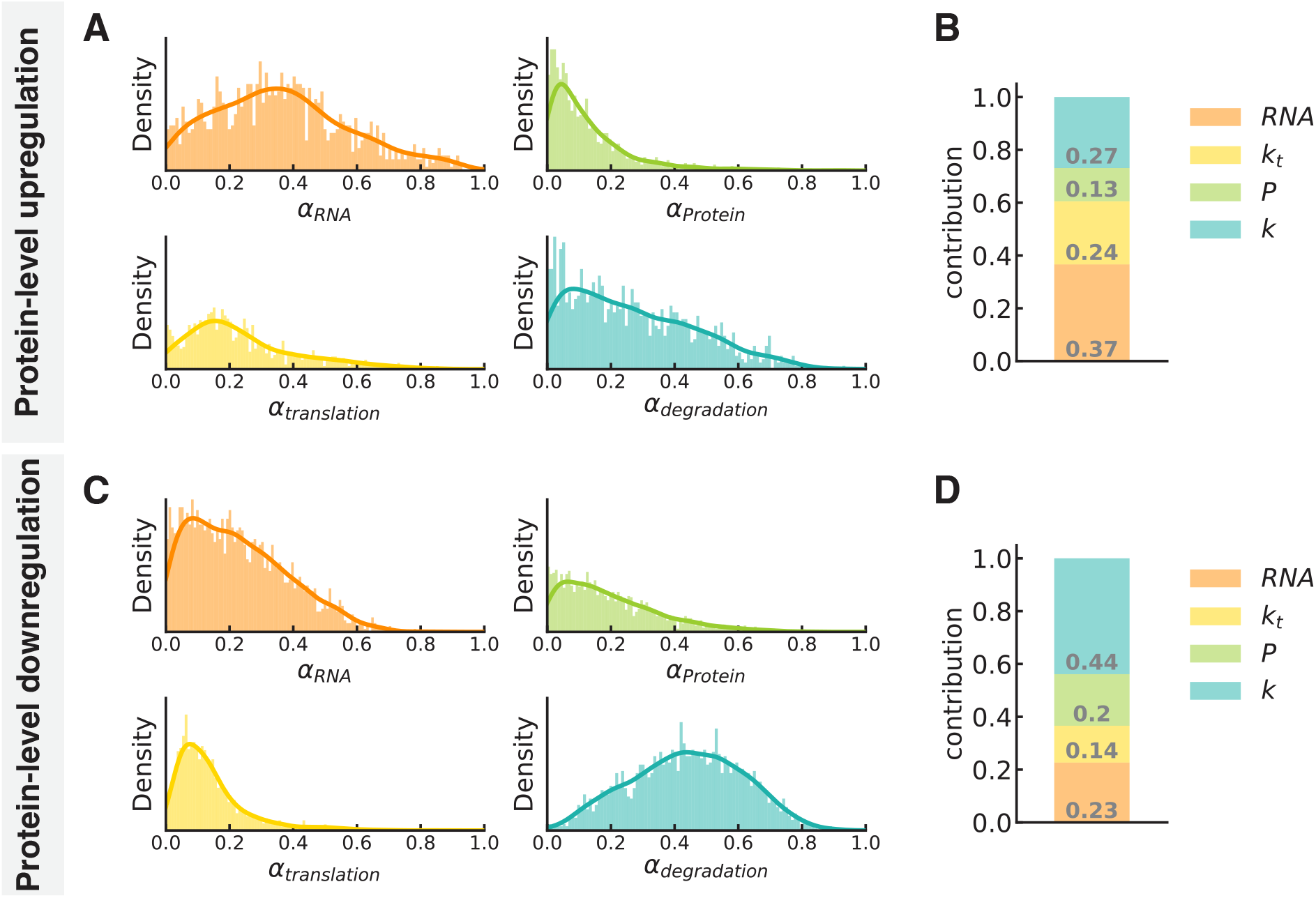
RNA level and protein degradation are the main contributor to protein flux changes for protein-level upregulation and downregulation, respectively. **A**. Distribution of the contributions of the different regulatory steps for the set of proteins exhibiting protein-level upregulation (*FI*(*P*) > 1). **B**. Barplot of the mean contributions of the different regulatory steps for the set of proteins exhibiting protein-level upregulation. **C**. Distribution of the contributions of the different regulatory steps for the set of proteins exhibiting protein-level downregulation (*FI*(*P*) < 1). **D**. Barplot of the mean contributions of the different regulatory steps for the set of proteins exhibiting protein-level downregulation. In **A**. and **C**., the plain lines represent the kernel density estimations from the associated histograms, for visualization purposes.

**Figure 1—figure supplement 3.**
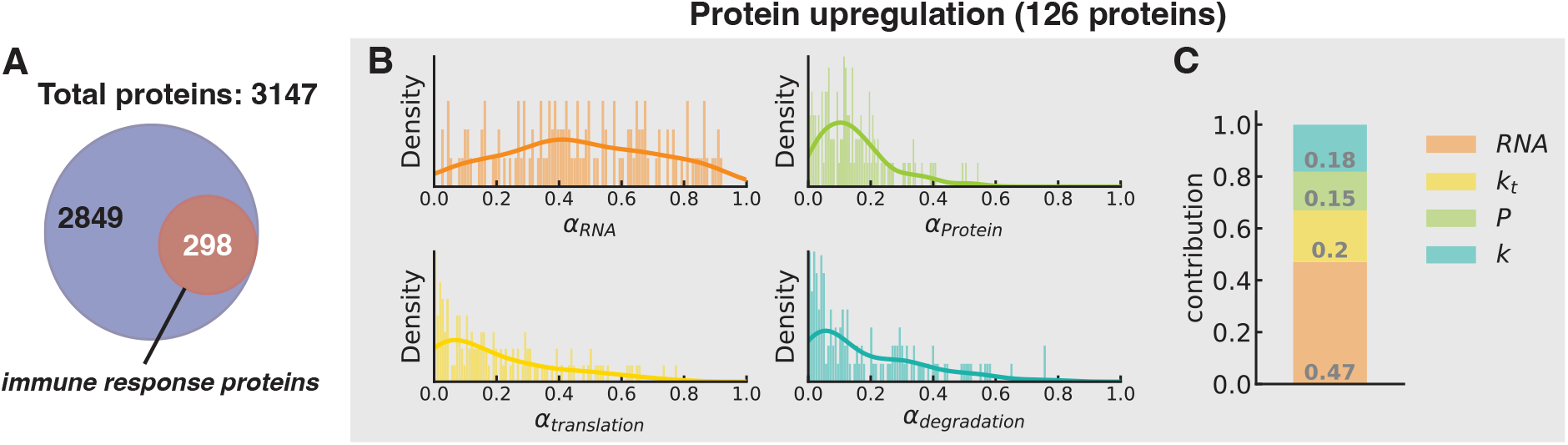
Change in protein flux for upregulated immune response proteins is mainly driven by RNA level changes. **A**. 298 immune response proteins were retrieved from Jovanovic et al. dataset using ImmuneDB (Rosenfeld et al., 2018). **B**. Distribution of the contributions of the different regulatory steps for the set of upregulated (*FI*(*P*) > 1) immune response proteins. The plain lines represent the kernel density estimations from the associated histograms, for visualization purposes. **C**. Barplot of the mean contributions of the different regulatory steps for the set of upregulated immune response proteins.

**Figure 1—figure supplement 4.**
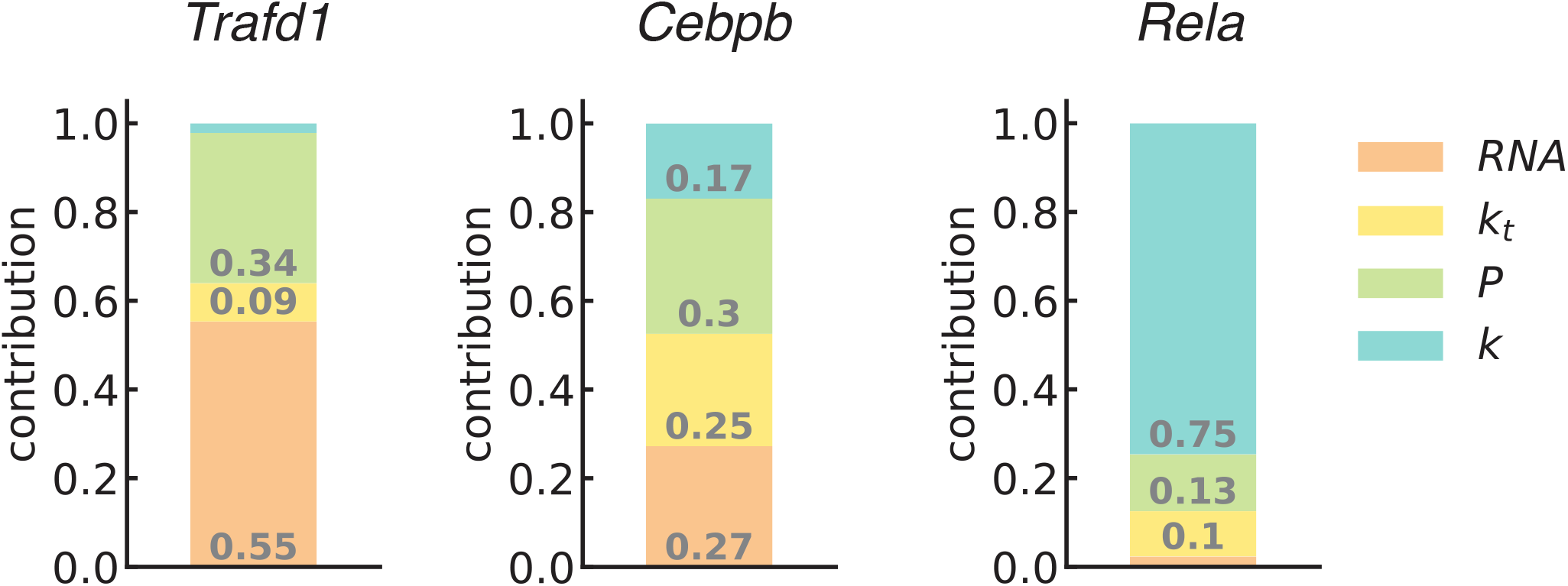
Immune response proteins exhibit variable contributions of the different regulatory steps. Barplot of the contributions of the different regulatory steps for the proteins Trafd1, Cebpb, and Rela.

**Figure 1—figure supplement 5.**
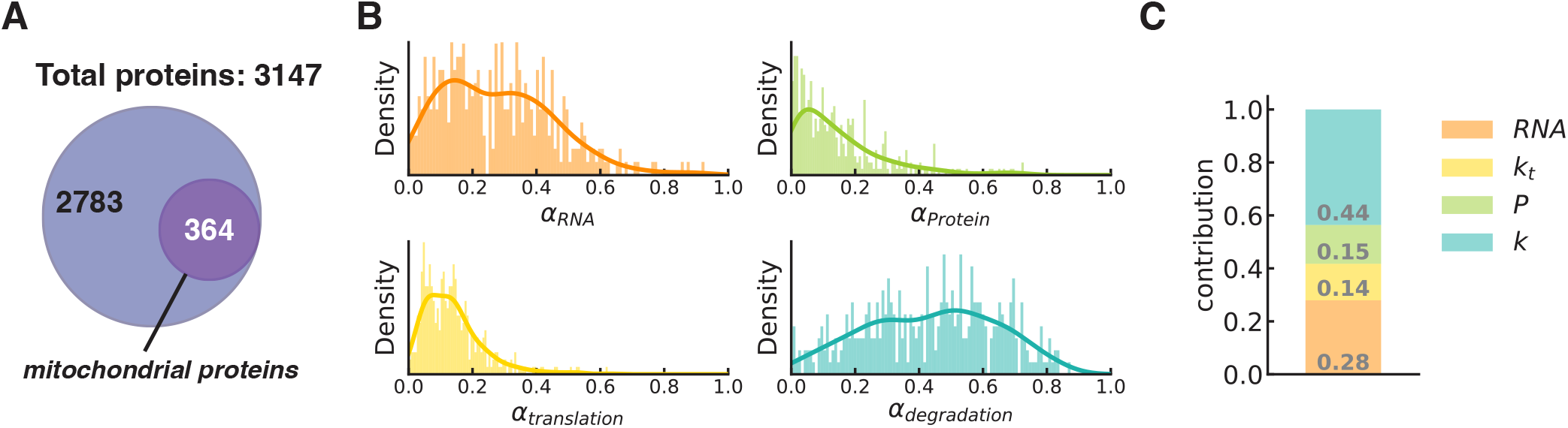
Change in protein flux for mitochondrial proteins is mainly driven by protein degradation. **A**. 364 mitochondrial proteins were retrieved from the dataset of Jovanovic et al. using MitoCarta 3.0 (Rath et al., 2021). **B**. Distribution of the contributions of the different regulatory steps for the set of mitochondrial proteins. The plain lines represent the kernel density estimations from the associated histograms, for visualization purposes. **C**. Barplot of the mean contributions of the different regulatory steps for the set of mitochondrial proteins.

**Figure 1—figure supplement 6.**
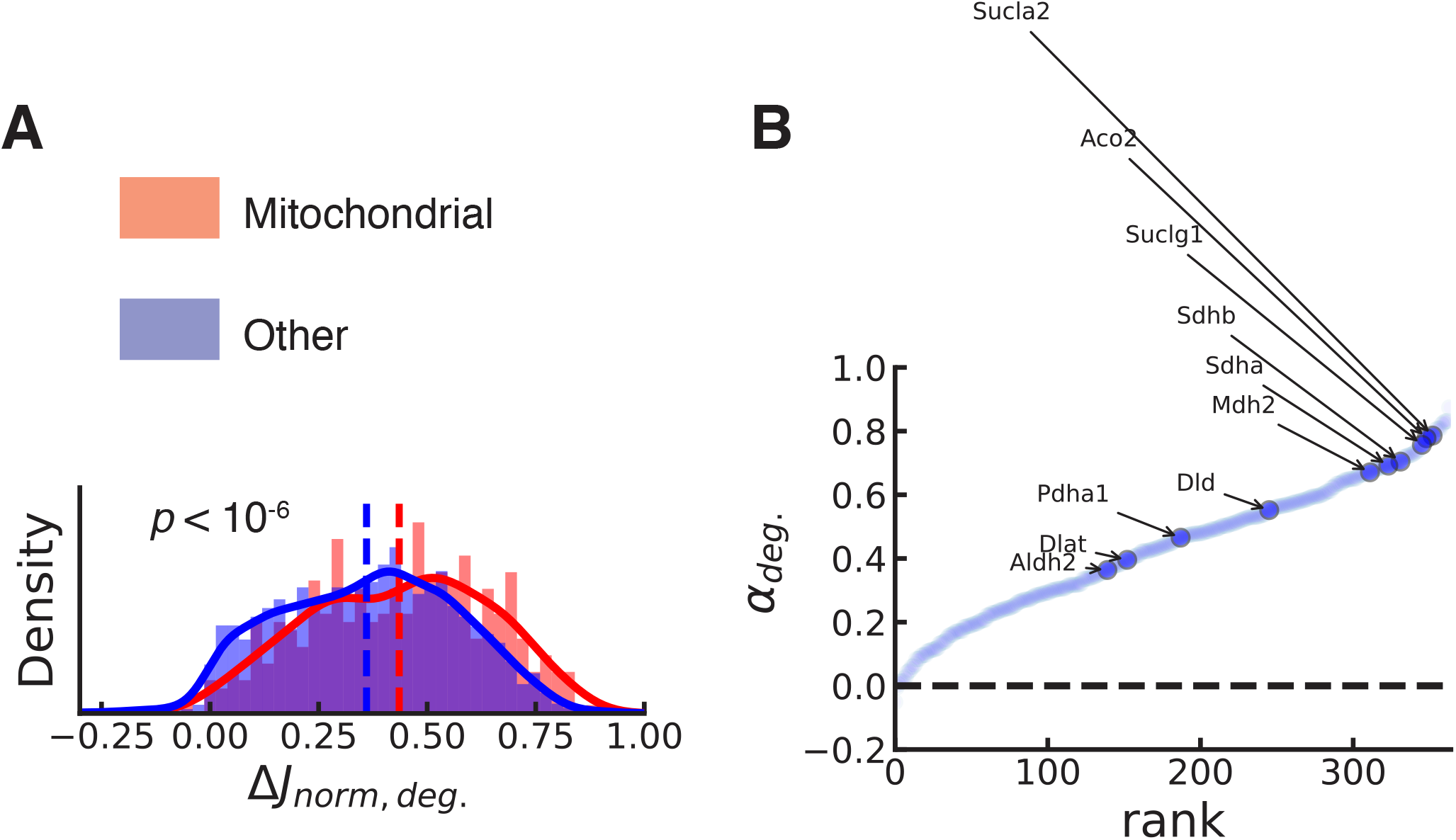
Increase in degradative flow of mitochondrial proteins leads to the metabolic rewiring of DCs after LPS stimulation. **A**. Distribution of the algebraic contribution of the protein degradation regulatory step *α*_*deg*_ for mitochondrial or non-mitochondrial (other) proteins. **B**. Ranking of mitochondrial proteins according to the algebraic contribution of the protein degradation regulatory step. Key mitochondrial enzymes of aerobic respiration are highlighted. The *P*-value was computed using Kolmogorov-Smirnov statistical test.

**Figure 1—figure supplement 7.**
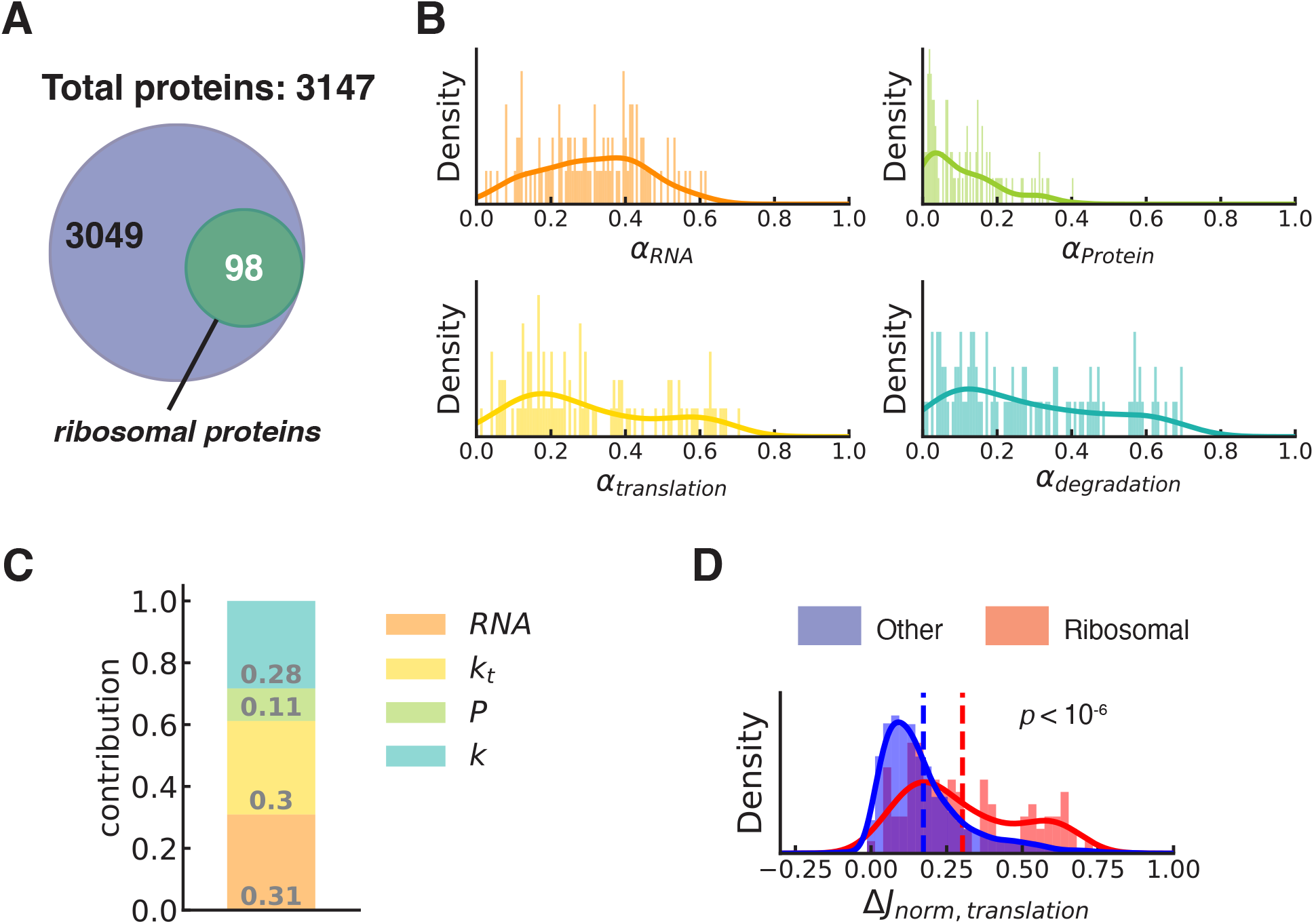
Translation is a key regulatory steps in protein flux changes of ribosomal proteins. **A**. 98 ribosomal proteins were retrieved from Jovanovic et al. dataset using the HGNC database (Tweedie et al., 2021). **B**. Distribution of the contributions of the different regulatory steps for the set of ribosomal proteins. **C**. Barplot of the mean contributions of the different regulatory steps for the set of ribosomal proteins. **D**. Distribution of the algebraic contribution *α*_*translation*_ for the sets of ribosomal or non-ribosomal (other) proteins. The plain lines represent the kernel density estimations from the associated histograms, for visualization purposes. *P*-value is computed using Kolmogorov-Smirnov statistical test.

**Figure 2—figure supplement 1.**
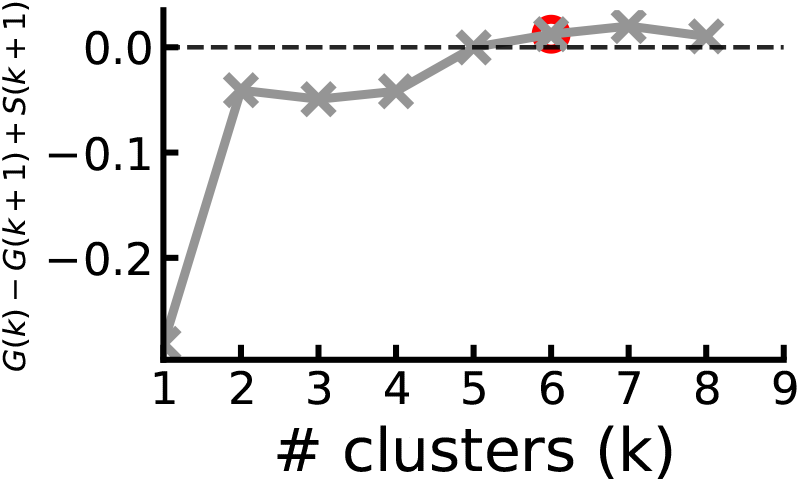
Gap statistics suggest the presence of six clusters in our dataset. Gap statistics were computed over 1000 replicates (Granger, 2022). The y-axis represents the “1-standard-error” style rule to select the optimal number of clusters. The optimal number of clusters is the smallest *k* such that *G*(*k*) ≥ *G*(*k* + 1) − *S*(*k* + 1) (Tibshirani et al., 2001). *G* is the Gap statistic and *S* is the simulation error (Tibshirani et al., 2001). The red dot represents our dataset’s optimal number of clusters with our clustering method (Methods and Materials).

**Figure 2—figure supplement 2.**
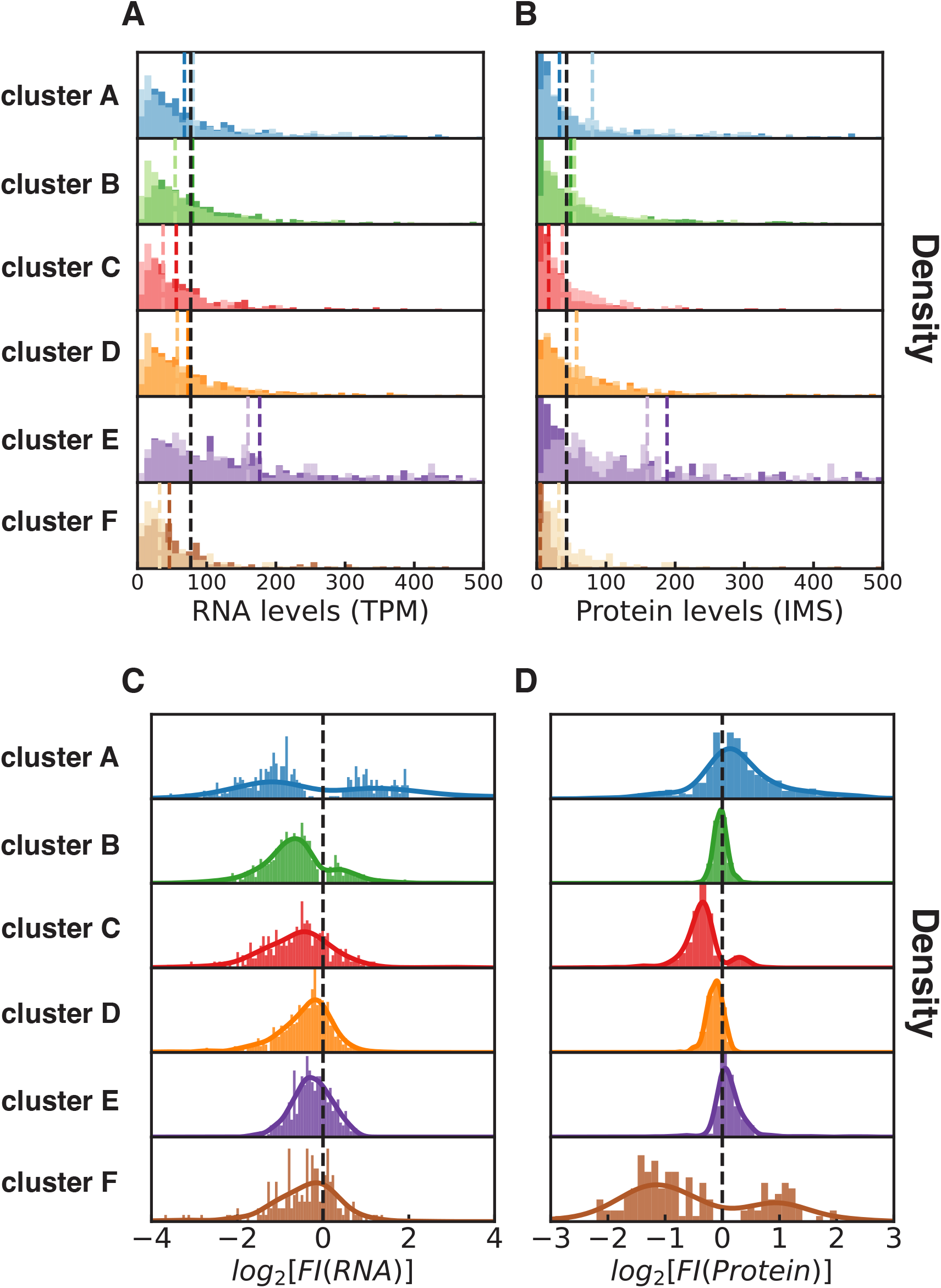
Regulatory modes are linked to versatile changes in protein flux, RNA, and protein levels. **A**. and **B**.: Initial (before LPS stimulation) and final (after 12h LPS stimulation: palish colors) distribution of RNA levels (**A**.) and protein levels (**B**.) for the six clusters of genes presented in ***Figure 2***. The black dotted line depicts the median over the whole protein dataset. The colored dotted line depicts the median of each respective cluster. **C**. and **D**.: Distribution of the LPS-mediated fold-increase in RNA levels (**C**.) and protein levels (**D**.), for the six clusters. The plain lines represent the kernel density estimations from the associated histograms,, for visualization purposes. The black dotted line indicates no changes between 0h and 12h LPS stimulation.

**Figure 2—figure supplement 3.**
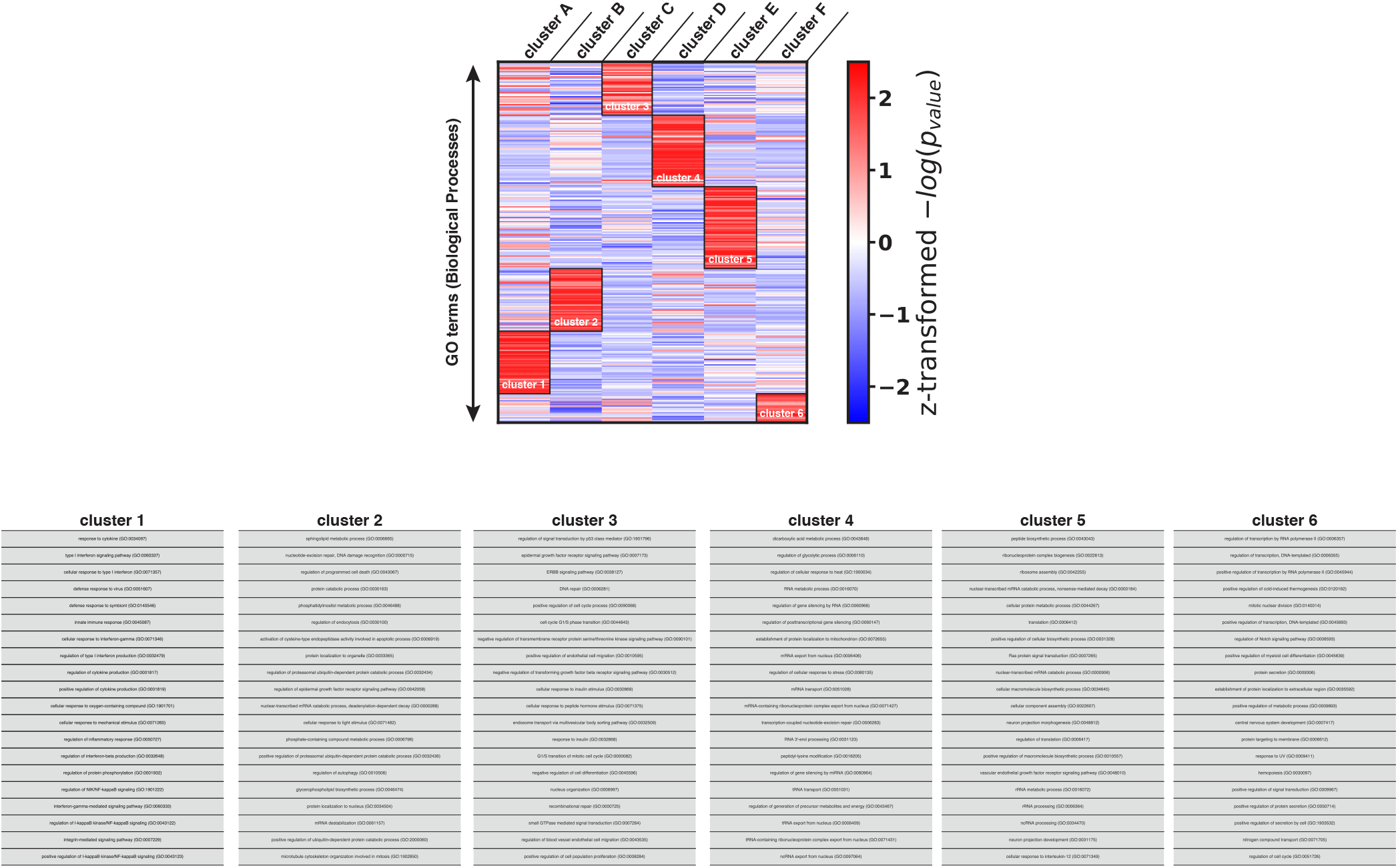
GO Biological processes terms can be clustered in six clusters of regulatory modalities. List of the 20 most statistically significant GO terms is given for each cluster. *log* is the *log*_10_ and *P*-value stands for the adjusted *P*-value.

